# The mitochondrial gene CMPK2 functions as a rheostat for macrophage homeostasis in inflammation

**DOI:** 10.1101/2021.11.01.466732

**Authors:** Prabhakar Arumugam, Meghna Chauhan, Thejaswitha Rajeev, Rahul Chakraborty, Deepthi Shankaran, Sivaprakash Ramalingam, Sheetal Gandotra, Vivek Rao

## Abstract

In addition to their role in cellular energy production, mitochondria are increasingly recognized as regulators of the innate immune response of phagocytes. Here, we demonstrate that altering expression levels of the mitochondria associated enzyme, cytidine monophosphate kinase 2 (CMPK2), disrupts mitochondrial physiology and significantly de-regulates the resting immune homeostasis of macrophages. Both CMPK2 silenced as well as constitutively over expressing macrophage lines portray mitochondrial stress with marked depolarization of their membrane potential, enhanced ROS and disturbed architecture culminating in the enhanced expression of the pro-inflammatory genes-IL1β, TNFα and IL8. Interestingly, the long-term modulation of CMPK2 expression resulted in an increased glycolytic flux of macrophages akin to the altered physiological state of activated M1 macrophages. While infection induced inflammation for restricting pathogens is regulated, our observation of a total dysregulation of basal inflammation by bidirectional alteration of CMPK2 expression, only highlights a critical role of this gene in mitochondria mediated control of inflammation.

## Introduction

Intracellular microbial infections compel host macrophages to activate selective cellular pathways that are targeted towards the elimination of the pathogen. In response to bacterial infections like Mtb, in addition to the expression of pro-inflammatory response mediators that facilitate the activation of bactericidal pathways (M1 polarized), recent evidence has demonstrated the robust activation of type I IFN response in macrophages(Galvan-Pena and O’Neill, 2014; Kumar et al., 2019; Manzanillo et al., 2012; Patrick and Watson, 2021). While the importance of the individual genes of this response cascade (interferon stimulated genes-ISG) in restricting viral nucleic acids is well recognized, its contribution in controlling bacterial infections is still unclear (Alphonse et al., 2021; Boxx and Cheng, 2016; Snell et al., 2017).

One of the early and highly expressed ISG in macrophages following infection is the mitochondria associated kinase CMPK2. CMPK2 is actively induced in response to several viral infections like HEV(Zhang et al., 2014), HIV(El-Diwany et al., 2018), Dengue virus (DENV)(Lai et al., 2021b), Nipah virus(Hassan et al., 2019), SVCV(Liu et al., 2019) and in response to LPS and poly IC(Kim et al., 2021) in macrophages. A recent study has demonstrated the involvement of the fish homolog-3nCmpk2 in antibacterial response by controlling bacterial colonization and protecting the intestinal barrier (Feng et al., 2021). In addition to its role in protective responses, CMPK2, has recently been implicated in activating host cell inflammasomes (Zhong et al., 2018) by virtue of its role in mitochondrial DNA synthesis thereby contributing to liver tissue injury and fostering premature atherosclerosis in SLE(Lai et al., 2021a; Luo et al., 2021).

This inherent duality of the inflammatory response: 1) leading to infection control (host beneficial) and 2) extensive tissue damage on uncontrolled activation (detrimental) signifies the need to effectively regulate the onset and end of the inflammation. Several mechanisms have been characterized for the control of inflammation by host phagocytes (Cohen and Mosser, 2013; Lawrence and Natoli, 2011; Sugimoto et al., 2016; Yan and Horng, 2020). Studies in the last decade have implicated a determinant connection between mitochondria and the inflammation balance of immune cells (Missiroli et al., 2020; Tiku et al., 2020; Wang et al., 2021). The enormous metabolic flexibility offered by mitochondria resident processes forms the basis of cellular plasticity facilitating fine-tuned responses to physiological conditions (Labbe et al., 2014; Picard and McEwen, 2018). The importance of mitochondrial health in terms of physiology, network architecture in immune cell function and response to infection is well recognized (Kikuchi et al., 1988; Rambold and Pearce, 2018; Weinberg et al., 2015). Activated M1 polarized macrophages demonstrate an important shift in the metabolic state relying completely on glycolysis for increased and faster production of energy(Blagih and Jones, 2012; Kikuchi *et al*., 1988; Newsholme et al., 1986; Rodriguez-Prados et al., 2010; Viola et al., 2019). This transition is associated with a distinctive change in mitochondrial structure and a diminished reliance on mitochondrial oxidative phosphorylation and increased production of mitochondrial ROS(Kelly and O’Neill, 2015; Labadarios et al., 1987; Mills et al., 2016; Van den Bossche et al., 2016). In line with our previous study on the robust induction of type I IFN response in Mtb infected macrophages (Shankaran et al., 2019), we show, that CMPK2, an ISG, is highly induced immediately following infection and is rapidly regulated to baseline levels in human macrophages. Altered expression of this gene is associated with a marked change in mitochondrial physiology indicating acute organellar stress; resting macrophages with constitutive silencing or over-expression of CMPK2 harbor significantly depolarized mitochondria with an atypical organellar network and heightened elaboration of mitochondrial ROS. This is reflected in the heightened levels of basal expression of inflammatory genes and a significant shift in macrophage metabolism to reliance on glycolysis resembling the M1 macrophages in these cells. Our study thus provides evidence for the importance of this gene in controlling inflammation in macrophages and the importance of regulated CMPK2 expression in protective responses, thereby highlighting an important link between mitochondrial metabolism and innate response kinetics of macrophages.

## Results

### Macrophages respond to infection/ TLR4 stimulation by inducing CMPK2

Given the early induction of type I IFN response in Mtb infections (Boxx and Cheng, 2016; Kovarik et al., 2016; McNab et al., 2015), we sought to investigate the importance of this pathway in the response dynamics of macrophages. The recent developments on the importance of mitochondrial physiology in host cell infection responses directed our attention to the mitochondria associated ISG-*CMPK2* that was highly induced following Mtb infection of macrophages. Infected THP1 macrophages showed a steady increase in *CMPK2* expression from 15-fold by 6h and attaining peak levels of ∼140 fold by 12h and thereafter maintained even at 24h of infection to greater than 100-fold (Fig. 1A). In line with previous reports (Zhong *et al*., 2018), we also observed nearly 20-fold increase in *CMPK2* expression in response to LPS stimulation while the other TLR ligands failed to alter this gene (Fig. 1B). This was also evident as a steady increase in CMPK2 protein levels in THP1 macrophages even at 48h of LPS stimulation (Fig. 1C). The complete loss of LPS induced expression in the presence of a TLR4 signaling inhibitor-CLI095, confirmed the dependence of *CMPK2* expression on TLR4 signaling in macrophages (Fig. 1D). Further, macrophages responded to *Salmonella enterica subsp. enterica* serovar Typhimurium (STM) infection, by early induction of *CMPK2* expression with 15-16 -fold increase by 3h wherein escalated further to ∼30 fold by 6h of infection (Fig. 1E) and this response was completely abolished with the addition of CLI095 confirming the TLR4 engagement dependent expression of this gene in macrophages (Fig. 1F). A decrease in expression by more than 50% was observed in the case of macrophages infected with Mtb and treated with CLI095 again supporting the importance of TLR4 signaling in *CMPK2* expression (Fig. S1).

**Fig. 1:**
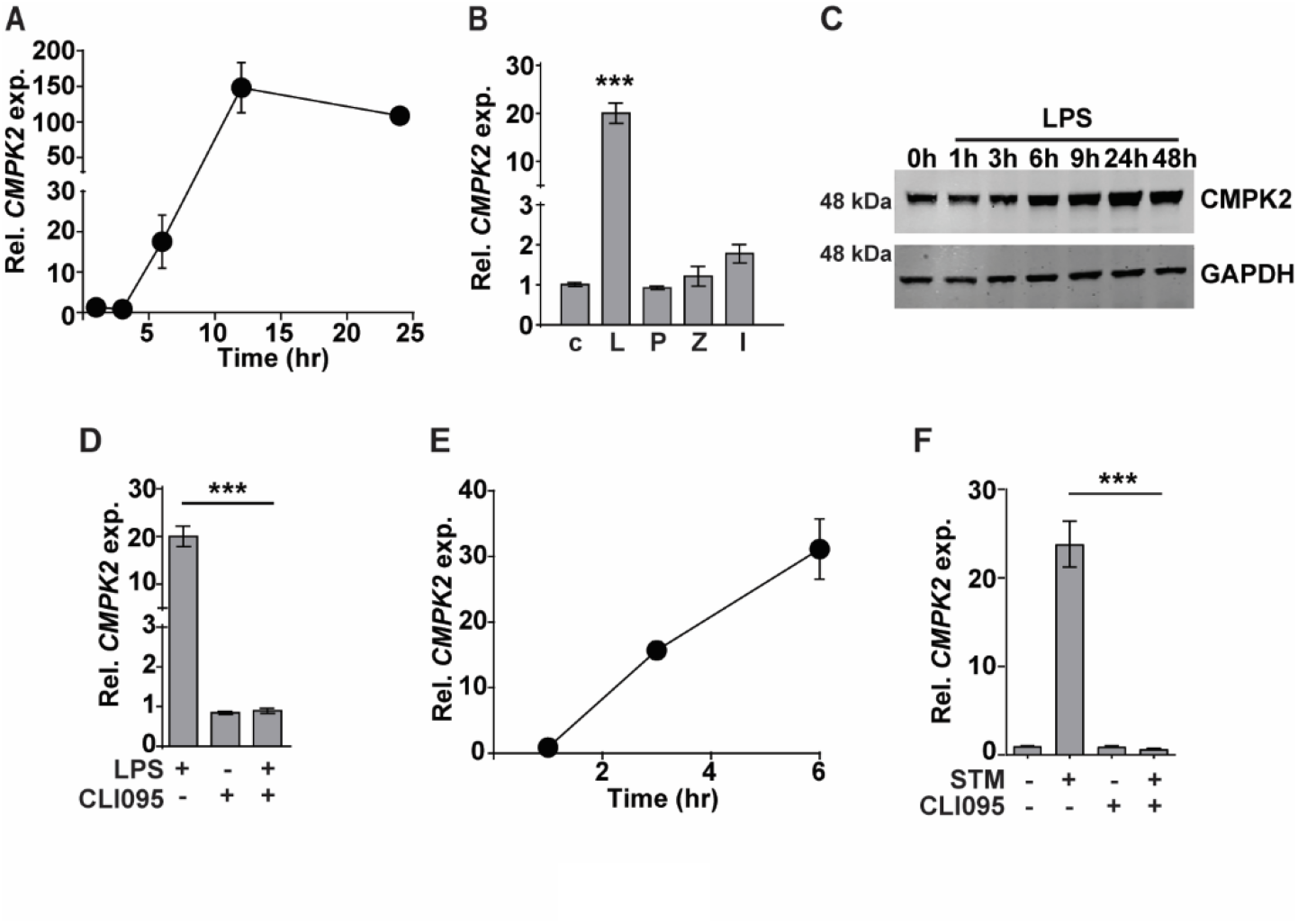
CMPK2 expression is induced following TLR4 stimulation of infection. [A-F] Analysis of CMPK2 expression in THP1 macrophages following: infection with Mtb at a MOI of 5 [A], stimulation with various TLR ligands-c: Untreated cells, L: LPS (10 ng/ml), P: Pam3CSK4 (20 ng/ml), Z: Zymosan (10 μg/ml), I: Poly IC (2 μg/ml) [B], LPS stimulation by immunoblotting with CMPK2-specific antibody [C], LPS stimulation with and without TLR4 inhibitor CLI095 [D], infection with STM [E], infection with STM in the presence of CLI095 [F]. Expression of *CMPK2* was quantitated by qPCR. Values are normalized with *GAPDH* and mean fold change compared to control cells + SEM from N=2/ 3 independent experiments.

### Modulation of CMPK2 activity affects the inflammation status of macrophages

To investigate the molecular function of CMPK2 in the macrophage response, we used *CMPK2* specific siRNAs (a-c) to specifically silence this gene in THP1 cells (KD) and achieved 60-75% decrease in gene expression in comparison to non-targeting siRNA (NT) (Fig. S2A). Only siRNA-c showed a significant decrease in protein levels in excess of 40% of the native CMPK2 expression and was used for further analysis (Fig. 2A). Given the critical role of proinflammatory cytokines in macrophage response to infection, we compared the levels of TNFα and IL1β in the NT and CMPK2 knockdown (KD) macrophages in response to Mtb infection. Mtb infection induced *TNFα* and *IL1β* expression by 200-500-fold in KD macrophages while these levels were 10-20-fold lower in the case of NT macrophages (Fig. 2B). Interestingly, expression levels of these genes were markedly elevated in the uninfected KD macrophages. This pattern was also reflected by the enhanced expression levels of other immune effectors like *IL8, IP10, VEGF* and decrease in *IL10* in naïve KD macrophages strongly alluding to an enhanced pro-inflammatory status of these cells even in the resting state (Fig. 2C). This profile of increased *TNFα* and *IL1β* gene expression was also observed in KD cells following LPS stimulation (Fig. S2B). This enhanced expression also correlated well with the increase in secreted TNFα and IL1β in the culture supernatant of LPS treated KD cells (Fig. 2D). Again, correlating with gene expression, naïve KD cells also secreted increased amounts of both TNFα and IL1β in the supernatants as compared to the non-detectable levels of the cytokine in naïve NT macrophages.

**Fig. 2:**
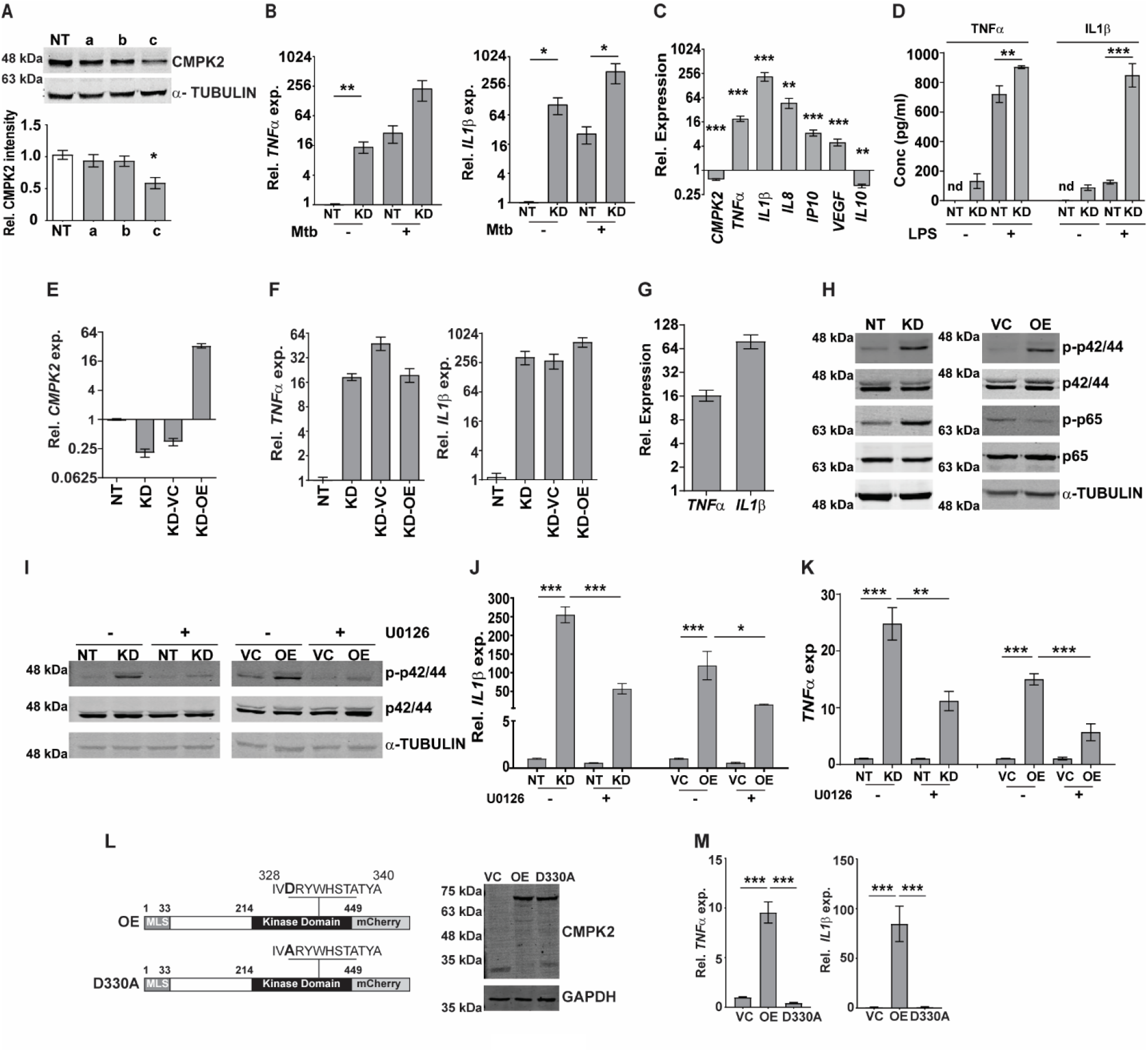
CMPK2 regulates the basal inflammation of macrophages. [A] Expression of CMPK2 in CMPK2 silenced macrophages by immunoblotting with specific antibodies. For immunoblotting, the expression of α-TUBULIN was used as control. One representative blot of 3 independent experiments is shown. The levels of CMPK2 expression are depicted as mean fold change + SEM of N=3 experiments by densitometry. [B] Expression of the *IL1β* and *TNFα* transcripts following infection with Mtb for 6h was evaluated in NT or KD macrophages by qPCR. [C] Basal level expression of transcripts *CMPK2* and cytokine genes in KD macrophages was estimated by qPCR with specific primers. Values are normalized with *GAPDH* and mean fold change compared to control cells + SEM from N=3 independent experiments. [D] Levels of TNFα and IL1β in the culture supernatants of THP1 stably expressing non-targeting (NT) or siRNA specific to CMPK2 (KD) estimated by ELISA. Values are represented as mean pg/ml of the cytokine in triplicate assays from N=3 experiments. [E] Expression of *CMPK2* in the CMPK2 silenced cells (KD) with stable expression of CMPK2 (KD-OE) or empty vector (KD-VC). The levels in the corresponding NT and KD cells are also shown. Values are represented as mean + SEM of the triplicate assays from N=3 experiments. [F] Expression of *TNFα* and *IL1β* in the CMPK2 silenced cells with stable expression of CMPK2 (KD-OE) or empty vector (KD-VC). The levels in the corresponding NT and KD cells are also shown. Values are represented as mean + SEM of the triplicate assays from N=3 experiments. [G] Expression of *TNFα* and *IL1β* in THP1 cells after stable expression of CMPK2. Values are represented as mean + SEM of the triplicate assays from N=3 experiments. [H] Analysis of activation of the ERK (p42/44) and NFκB (p65) signaling pathways in NT, KD, VC and OE macrophages by immunoblotting with antibodies specific for the phosphorylated (active) and non-phosphorylated forms of the proteins. Representative blot of one experiment out of 3 individual assays is shown. Expression of α-TUBULIN was used as control. [I] ERK phosphorylation in macrophages with and without treatment with ERK specific inhibitor U0126. Antibodies specific for the phosphorylated (active) and non-phosphorylated forms of the proteins were used to probe cell extracts. Expression of α-TUBULIN was used as control. Blots are representative of two independent experiments. [J-K] Expression of *IL1β* [J] and *TNFα* [K] in the 4 types of macrophages after treatment with U0126 was analyzed by qPCR. Cells left untreated were used as control. The relative gene expression folds in triplicate assay wells are represented with respect to *GAPDH* as mean + SEM for N=2. [L] The schematic of mutated catalytic site also depicted (D330A). Immunoblot analysis of CMPK2-mCherry fusion protein using an antibody specific for mCherry protein in protein lysates of VC, OE and D330A THP1 cells with mCherry specific antibody. GAPDH was used as control. [M] Expression of *TNFα* and *IL1β* in THP1 cells after stable expression of vector alone (VC), full length CMPK2 (OE) and catalytic mutant of CMPK2 (D330A). The relative gene expression folds in triplicate assay wells are represented with respect to *GAPDH* as mean + SEM from N=4.

Given the inflammatory phenotype observed with decreased CMPK2 expression in THP1 macrophages, we attempted to reverse this by complementing a functional copy of the gene in the silenced cells. We expressed the complete CMPK2 as a mCherry fusion protein in pCDNA3.1 and analyzed the basal expression levels of *IL1β* and *TNFα*. As expected, *CMPK2* expression was ∼30 folds higher than in the empty vector control macrophages (Fig. 2E). Contrary to our expectations, this enhanced expression of the *CMPK2* could not reverse the alteration in basal inflammation of these cells: nearly similar levels of *IL1β* and *TNFα* were observed in the complemented cells as in the silenced cells (Fig. 2F). We hypothesized that similar to the situation with CMPK2 silenced cells, an increase in expression of CMPK2 was also detrimental to the inflammation homeostasis of macrophages. To test this, we analyzed the pro-inflammatory status of wild type (Wt) THP1 cells stably expressing CMPK2-mCherry. Enhanced expression of *CMPK2* transcript in excess of 100-fold basal levels was observed in the over expressing (OE) cells compared to the vector control (VC) cells (Fig. S2C). The fusion protein of CMPK2 and mCherry was also clearly detected in protein extracts of the OE strains by immunoblotting (Fig. S2D). Again, this enhanced expression of CMPK2 resulted in heightened levels of *TNFα* (12-15-fold) and *IL1β* (60-80-fold) in OE cells (Fig. 2G).

In an effort to understand the basis for the hyper-inflammatory state of KD cells, we compared the immune signaling cascades of these cells with the NT cells in the basal state. Consistently, we observed enhanced activation of ERK and NFκB with 3-fold higher levels of phospho-p42/44 (ERK) and p65 NFκB (Fig. 2H, S2E) without any change in phospho-JNK/SAPK and p38 MAP kinase between the two cells (Fig. S2F). This pattern of enhanced ERK phosphorylation was also observed in the case of OE cells in comparison to VC cells. The use of an ERK signaling specific inhibitor U0126 not only reduced the levels of phospho-p42/44 in both the KD and OE cells significantly (Fig. 2I), but also significantly reduced the elevated levels of both *IL1β* and *TNFα* in both these cell types, further confirming the importance of ERK signaling in mediating the hyper inflammatory phenotype of these macrophages (Fig. 2J, 2K).

Human mitochondrial CMPK2 has been shown to be a nucleotide kinase involved in mitochondrial DNA synthesis (Fig. 2L). In an effort to probe the importance of the kinase domain, we substituted the active site aspartate to alanine (D330A) as a kinase dead variant of CMPK2 (D330A) and analyzed the functional consequence of this expression on cytokine gene expression. Despite the strong and stable expression of D330A, comparable to the Wt, CMPK2 (OE), both at the transcript (Fig. S2G) and protein (Fig. 2L) level, there was no evidence of increased basal inflammation. Contrasting with the significantly elevated levels of *TNFα* and *IL1β* in the OE cells (∼10 and ∼80 fold, respectively over basal levels), THP1 cells transfected with D330A, harbored normal levels of gene expression (Fig. 2M) emphasizing the importance of the kinase activity of CMPK2 in regulating basal inflammation in macrophages.

### Increased inflammation of KD macrophages is associated with altered mitochondria and augmented ROS

We presumed that identifying the precise localization of CMPK2 would reveal clues to its molecular role in inflammation and mitochondrial membrane dynamics. We first validated the mitochondrial localization of CMPK2 by immunoblotting subcellular fractions. With previous reports indicative of the mitochondrial localization of CMPK2 (Xu et al., 2008) and our observation of a majority of the protein in the mitochondrial fraction (Fig. 3A), we tested purified mitochondrial fractions to pinpoint its location by standard biochemical and immunoblotting analyses. While treatment with Proteinase K (removes the surface exposed proteins), completely removed the outer membrane protein TOM20, nearly 30% of CMPK2 was present in the pellet fraction of intact mitochondria, similar to that observed for the other mitochondrial membrane associated protein – TIM50 (Fig. 3B) suggesting its association with the outer membrane of mitochondria. Most of the matrix protein-SOD2 remained in the pellet and was lost only with complete solubilization with Triton X-100. The peripheral association of CMPK2 with the mitochondrial membrane was also revealed by a relatively higher amount of CMPK2 in the soluble fraction of mitochondria treated with Na_2_CO_3_ in contrast with the integral membrane protein-TOM20 (Fig. 3C). We hypothesized that two distinct possibilities could account for the enhanced basal levels of inflammation observed in the CMPK2 expression altered cells (KD and OE) - 1) enhanced expression at all times or 2) a temporal activation of the signaling with delayed decay kinetics. To test this, we checked the expression of *IL1β* in the two cell lines during the monocyte to macrophage transition upon PMA treatment. While the monocytes did not show any difference in *IL1β* expression, activation with PMA induced distinct profiles of *IL1β* activation in the control (NT) or CMPK2 silenced (KD) cells. A rapid increase in *IL1β* within a day of activation (∼700 fold) was followed by a steady decline by day 3 of treatment further reaching basal levels by day 4 in the control cells (Fig. 3D). CMPK2 silenced macrophages, however, displayed a protracted response kinetics with a slower rise in expression levels ∼1.5-2 folds lower than control cells at 24 h of activation with PMA. Between 24h and 72h of treatment, the levels were 3-5 fold higher in KD with a delayed decline of the levels to ∼500 fold by day 4 and did not fall to basal levels even by day 5. Macrophage inflammation is strongly associated with ROS activation. To check this, we compared the levels of mitochondrial ROS in NT and KD macrophages by using a ROS specific stain-MitoSOX Red. Mitotracker deep red was used as control for the levels of mitochondria in the two cell types. While similar levels of mitotracker deep red staining hinted at unaltered mitochondrial levels, the level of ROS (mitosox) was nearly 1.5-fold higher in KD cells as compared to control NT macrophages (Fig. 3E). The importance of mito-ROS in enhanced inflammation in KD cells was further evident when the cells were treated with a specific inhibitor-mitoquinone (MQ). Addition of MQ early (immediately after PMA treatment) significantly reduced the expression of *IL1β* in these cells (Fig. 3F); use of MQ at later stages of differentiation (48h and 72h after PMA addition) did not alter the inflammation in the KD cells (Fig. S3A, B). Mitochondrial membrane polarization is a critical component of organelle integrity and physiology with alterations in membrane potential leading to ROS in macrophages (Suski et al., 2012; Vyssokikh et al., 2020). To test if the ROS was associated with change in membrane potential, we compared the extent of TMRE staining (a dye that shows a potential dependent differential partitioning across mitochondrial membranes) in cells. As seen in Fig. 3G, the mitochondria of both KD and OE macrophages exhibited significantly lower staining with TMRE, similar to the NT/VC cells treated with the proton pump uncoupler-CCCP, suggestive of a loss of membrane integrity in the cells even at the basal level.

**Fig. 3:**
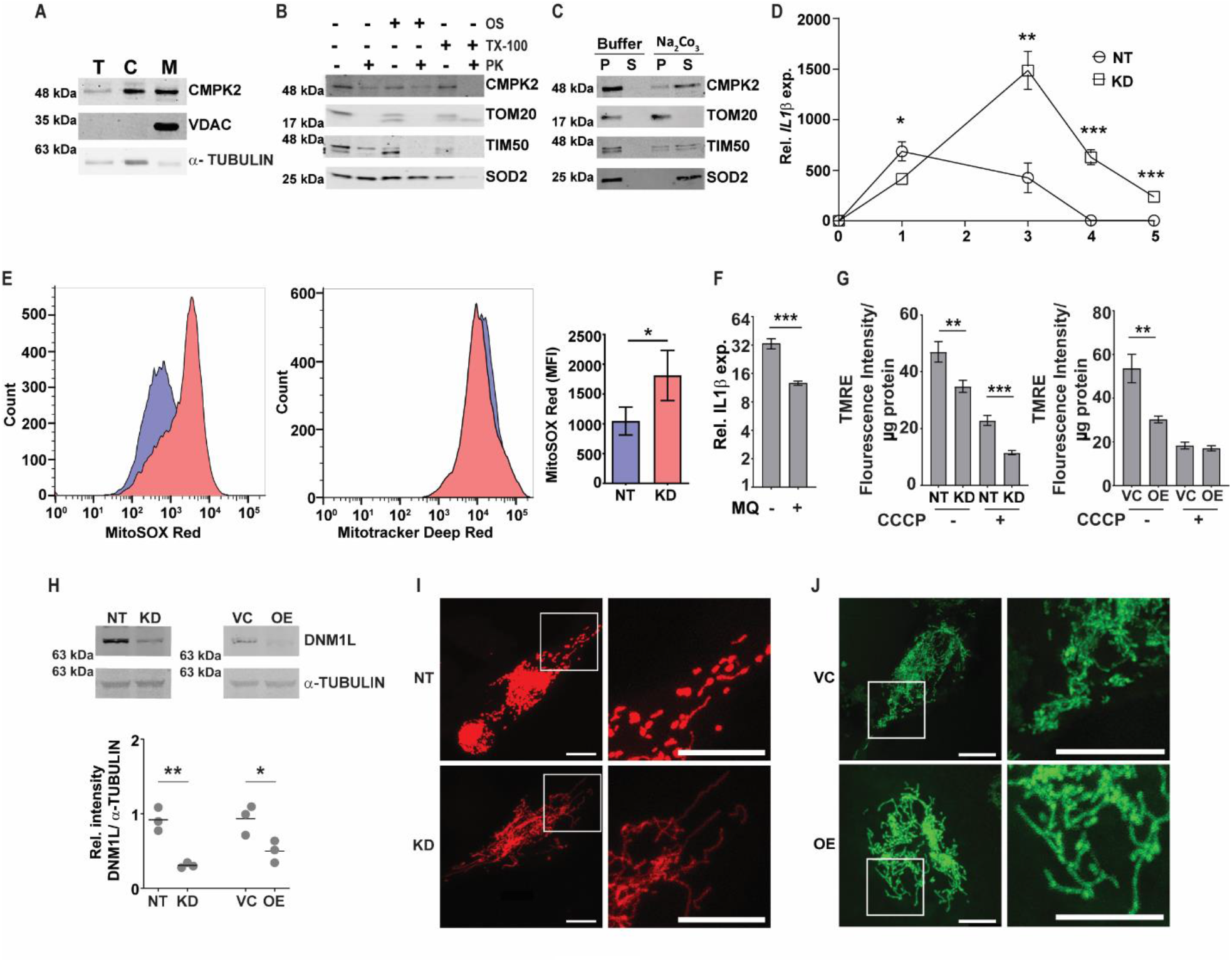
Modulation of CMPK2 affects the mitochondrial physiology in macrophages. [A] Expression of CMPK2 in subcellular fractions of THP1 macrophages by immunoblotting with specific antibodies, T-total cell extract, C-cytoplasmic fraction and M-Mitochondria. Expression of cytosolic protein α-TUBULIN and mitochondria resident protein VDAC are also represented. Blots are representative of three independent experiments. [B] Mitochondrial fraction was subjected to Proteinase K treatment in the presence or absence of Triton X-100 and given osmotic shock (OS) and analyzed by immunoblotting. Data are representative of three independent experiments. [C] Mitochondrial fraction was incubated with mitochondrial buffer with and without Na2CO3 and centrifuged at 13,000 rpm for 15 min. The pellet (P) and supernatant (S) fractions were immunoblotted. Data are representative of three independent experiments. [D] Kinetic profile of IL1β expression in NT and KD during differentiation of monocytes to macrophages. Expression was checked at different time intervals after PMA treatment. Values are mean fold change in expression with respect to GAPDH + SEM triplicate assays of N=2 experiments. [E] Analysis of mitochondrial ROS in NT or KD macrophages. Cells were stained with MitoTracker Deep Red (as an internal control) and ROS specific MitoSOX Red and specific populations were quantified by FACS. The histogram plots of a representative experiment of N=4 is depicted. The extent of MitoSOX red mean fluorescence intensity (MFI) + SEM is represented graphically in the inset. [F] Expression of IL1β in the macrophages with the addition of a specific mitochondrial ROS inhibitor-MQ (1 μM) at day1 of PMA treatment was analyzed by qPCR. Values are mean fold change in expression with respect to GAPDH + SEM for triplicate assays of N=3 experiments. [G] Analysis of mitochondrial membrane potential in NT, KD, VC and OE macrophages by TMRE staining. Fluorescence values normalized to protein in samples are represented as mean fluorescent intensity + SEM for triplicate assays of N=3 experiments. [H] Expression of DNM1L by immunoblotting with specific antibody was analyzed and is depicted along with the levels of α-TUBULIN as a control. Relative intensity values are depicted as mean+ SEM of N=3. [I, J] Mitochondrial architecture in control (NT) and KD [I] or VC and OE [J] macrophages were analyzed by confocal microscopy. A representative image is depicted with the scale bar representing 10μm, the region shown in higher magnification is depicted with the box.

Given the importance of mitochondrial fission-fusion dynamics as a stress mitigating factor we tested if alteration in CMPK2 expression and the associated stress resulted in aberrant mitochondrial replication dynamics. In line with the previous report (Youle and van der Bliek, 2012) both KD and OE cells showed decreased levels of DNM1L (crucial for mitochondrial fission) in comparison to NT and VC cells, respectively, (Fig. 3H), contrasting with an unaltered expression of the mitochondrial fusion associated proteins (MFN1 and OPA1) in these cells (Fig. S3C, S3D). This was further evident in the larger and more dense mitochondrial network of these organelles in the KD and OE cells as compared to the small uniformly distributed mitochondria in the control cells (Fig. 3I, 3J).

### Regulated levels of CMPK2 are critical for the normal metabolic activity of macrophages

In an attempt to understand the basis for this dysregulated inflammatory gene expression in both the KD and OE macrophages, we performed global sequencing of transcripts and compared the expression profiles with the control macrophages -NT and VC, respectively. Strikingly, again, on comparison of the differentially expressed genes, inflammatory cytokine genes like *IL1β, IL8* and *TNFα* were commonly upregulated in both the KD and OE macrophages (Fig. 4A). This consistency in gene expression profiles was also evident with the significantly large numbers of genes increased (515) or decreased (500) in macrophages with abnormal CMPK2 expression (Fig. 4A inset). A significant deregulation of basal level inflammation was also evident in Gene Set Enrichment Analysis (GSEA) with heightened inflammation response as a key gene family common to the two macrophage lines (Fig. 4B). Interestingly, the hypoxia response was highlighted as one of the highly upregulated gene family that was observed in both the KD and OE macrophages. Most of the genes involved in the response were upregulated anywhere between 2 and 16 folds in these cells (Fig. 4C). With hypoxia primarily regulated by HIF1α in cells, we probed the levels of this protein in extracts of these macrophages. HIF1α levels were elevated ∼2 folds in the OE cells as compared to the control VC cells, similarly higher level of HIF1α was also observed in extracts of the KD cells (Fig. 4D).

**Fig. 4:**
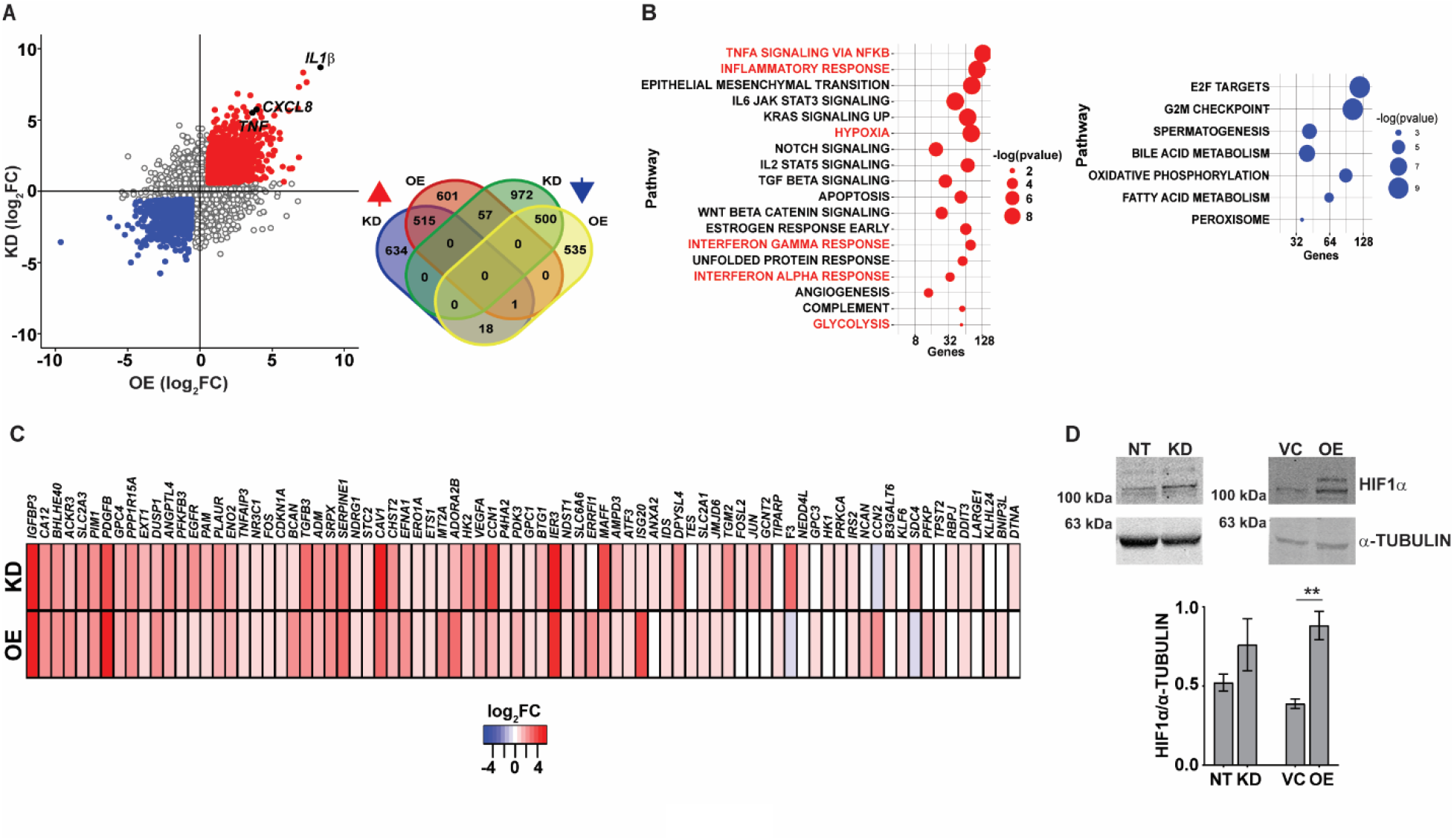
Macrophages with dysregulated CMPK2 display increased hypoxia. [A] Scatter plot of genes differentially expressed in KD or OE macrophages relative to the expression levels in the control macrophages (NT or VC), respectively. Change in expression is depicted as log_2_ fold change in expression. The number of genes up- and down-regulated in the macrophages are depicted as a Venn diagram (inset). [B] GSEA hallmark pathway enrichment analysis of the commonly up and down regulated genes in the KD and OE macrophages are represented as a bubble plot. X axis is the number of genes of the pathway and size of the bubble depicts significance (-log P value). [C] Heat map of the hypoxia response gene expression in the CMPK2 dysregulated macrophages KD and OE. The values represent log_2_ fold change from the corresponding control cells. [D] Analysis of HIF1α in protein extracts of NT, KD, VC and OE cells. Equal amounts of protein from the different extracts were probed with specific antibodies, a representative blot of 3 independent experiments is depicted. The change in expression level of HIF1α with respect to α-TUBULIN was calculated by densitometric analysis and is graphically represented.

### Macrophages with dysregulated CMPK2 expression levels portray signatures of activated M1 cells

The increased hypoxia, mitochondrial ROS and inflammation observed in the KD and OE cells were reminiscent of M1 activated macrophages. To validate this, we analyzed the expression profiles of previously identified signatures of M1 and M2 polarized macrophages in our gene sets (Martinez et al., 2006). In agreement with our hypothesis, both the KD and OE macrophages showed an increase in the expression of genes of M1 macrophages with a majority of the M2 specific signatures either unchanged or with reduced expression levels (Fig. 5A). Further validation of a M1 bias was visible in the physiological state of the cells. Analysis of respiration rates of cells revealed a distinct pattern with a considerable decrease in the oxygen consumption rates of macrophages with altered expression of CMPK2. Both the KD and OE macrophages displayed 2-2.5 folds lower oxygen consumption with delayed kinetics as opposed to the control macrophages NT and VC (Fig. 5B) implicating a strong shift of macrophages with altered CMPK2 expression towards a pro-inflammatory phenotype and the important role for CMPK2 in maintaining homeostasis of macrophages. Further, both these cell types harbored increased expression of genes involved in glycolysis (Fig. S4A, 5C) that manifested as a significant increase in the levels of glycolysis intermediates like fructose bis-phosphates, DHAP, culminating in markedly high levels of lactate (Fig. 5C). Recent studies have demonstrated a strong link between serine biosynthesis/ metabolism to the inflammatory profiles and mitochondrial physiology of cells (Kurita et al., 2021; Wang et al., 2020; Yu et al., 2019). Strikingly, the serine-glycine biosynthetic pathway was one of the dominant metabolic axes downregulated in these 2 cell types in comparison with the control cells (Fig. S4B). The expression levels of the key enzymes involved in this pathway was validated by qRTPCR. Consistently, in both the KD and OE cells, expression levels of *PHGDH, PSAT1* and *PSMT1* were significantly lower in comparison to the control NT and VC cells (Fig. 5C). This decrease in expression levels is reflected as ∼2-fold decrease at the metabolite level of serine in the KD and OE cells. Given an important role for PHGDH in controlling inflammation under hypoxic conditions(Huang et al., 2021; Wilson et al., 2020) and our observation of a significant decrease in expression levels combined with hypoxic conditions and heightened pro-inflammatory gene expression in the KD and OE cells, we hypothesized an important role for this axis in the observed phenotypes in this study. To investigate this, we specifically inhibited PHGDH in THP1 Wt cells by the specific inhibitor-NCT-503. As a control, cells were treated with an inactive inhibitor of PHGDH and evaluated the expression profiles of *IL1β* and *TNFα* in these cells. Use of the inactive inhibitor did not alter the expression from the untreated cells, in contrast to the significant increase in transcript levels by ∼5 and 2.5 folds for these genes with the inhibitor, substantiating our supposition (Fig. 5D).

**Fig. 5:**
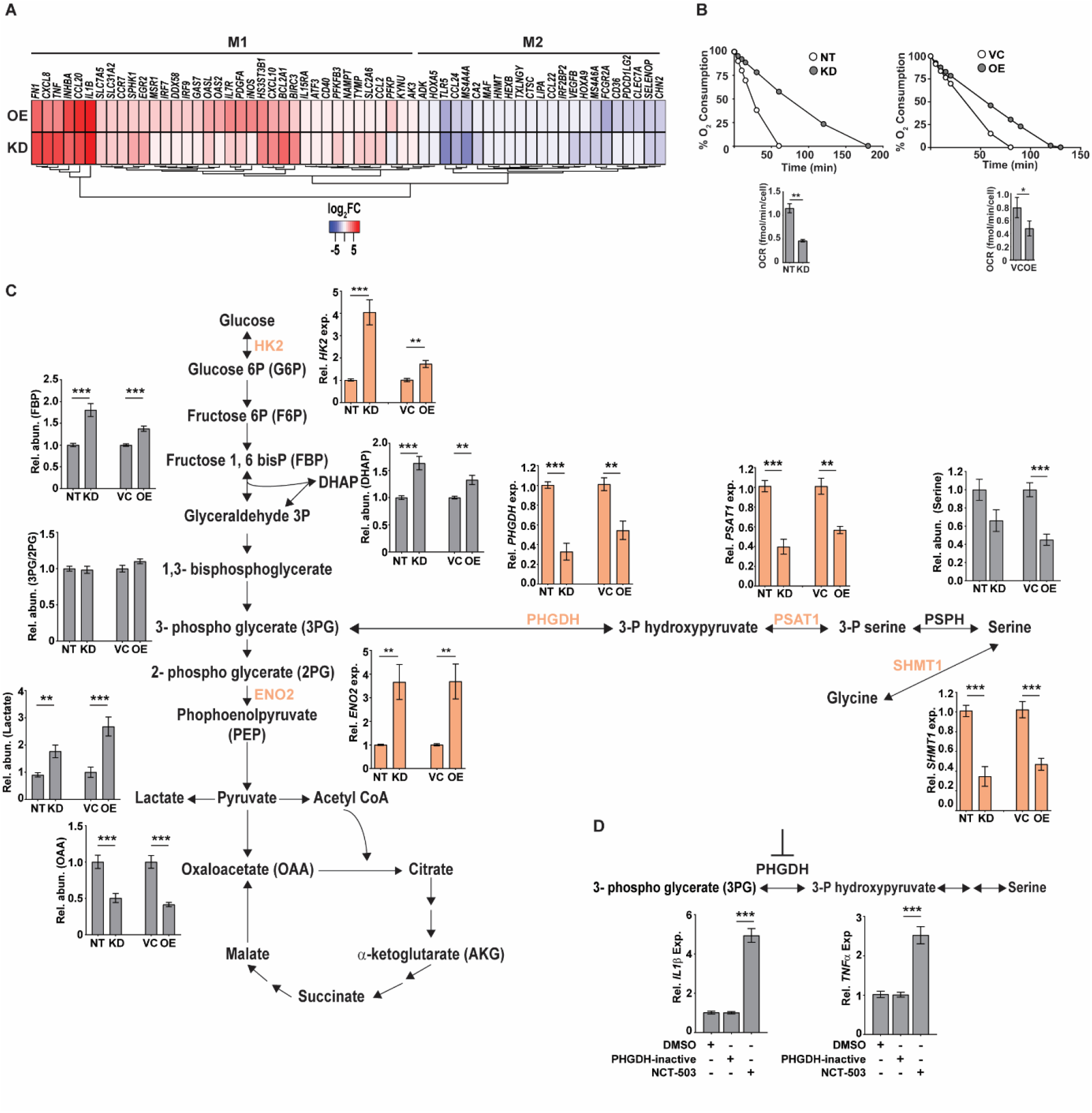
Macrophages with dysregulated CMPK2 are metabolically similar to activated M1 macrophages with higher glycolytic flux. [A] The expression patterns of genes specific to M1 and M2 activated macrophages in the CMPK2 dysregulated macrophages KD and OE are represented as heat maps. The values represent log_2_ fold change from the corresponding control cells. [B] Continual estimation of oxygen consumption in NT, KD and VC, OE macrophages until 3h with Oroboros oxygraph. The level of oxygen consumption was calculated and mean OCR + SEM of N=3 experiments is shown. [C] Metabolite levels in the CMPK2 silenced (KD) and over-expression (OE) and their respective control macrophages were assayed from cellular extracts by MS. The levels of individual metabolites in the KD and OE macrophages are represented as relative abundance compared to control cells as mean + SEM from 2/3 independent experiments (N=3). Expression of genes involved in the glycolysis and serine to glycine biosynthetic pathway estimated by qRT-PCR was also shown. [D] Expression of *IL1β* and *TNFα* in THP1 cells treated with the serine biosynthesis inhibitor-NCT-503 and its inactive form-PHGDH-inactive. Expression values are mean fold change in expression with respect to *GAPDH* + SEM triplicate assays of N=2 experiments after 6h of treatment.

### CMPK2 plays a critical role in the macrophage ability to control infection

To understand the impact of altered CMPK2 expression on macrophage inflammatory properties, we evaluated its ability to control the intracellular growth of Mtb. We observed a 2-3 fold decrease in intracellular bacterial numbers in these macrophages as compared to NT late in infection (day 5), KD cells were able to control bacterial levels to input levels by this timepoint compared to growth of about 5-6 folds in the NT macrophages (Fig. 6A). This pattern of increased bacterial control was also evident in STM infected KD macrophages with 2-3 fold lower bacterial numbers after 24 h of infection (Fig. 6B).

**Fig. 6:**
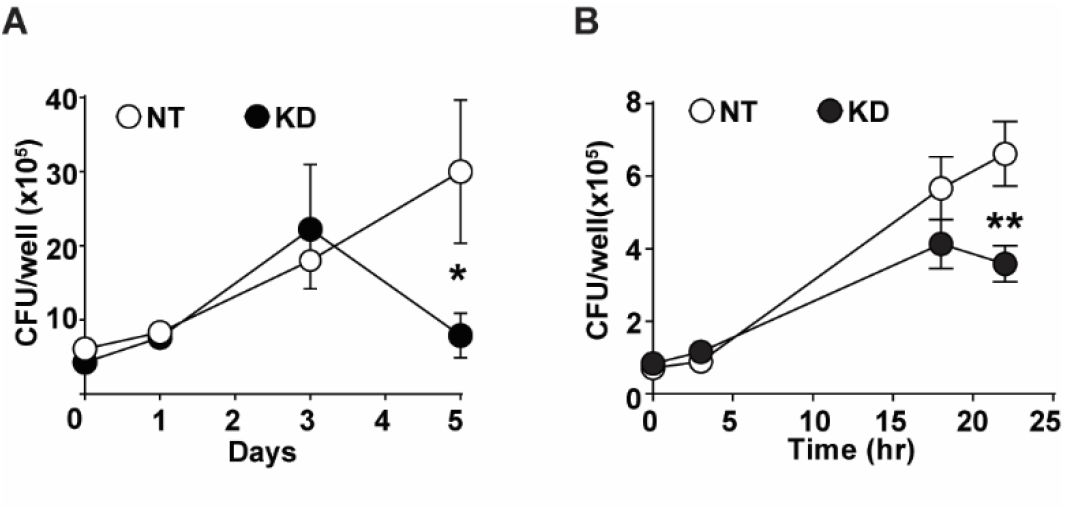
CMPK2 regulates the bactericidal activity of macrophages. [A] Growth kinetics of Mtb in NT and KD macrophages at different times post infection at a MOI of 5 for 6h. Values are mean CFU + SEM values in triplicate assays of N=4. [B] Growth kinetics of STM in NT and KD macrophages at different times post infection at a MOI of 10 for 20 min. Values are mean CFU + SEM values in triplicate assays of N=4.

## Discussion

Macrophages, the primary cells that sense and respond to any infectious agents or tissue damage, are critical components of vertebrate innate immune responses. Being endowed with the capacity to initiate and manifest strong infection control programs to a variety of pathogens, they often dictate both acute and chronic outcomes of infection in the host (Gong et al., 2020; Mosser et al., 2021). Recent evidences also hint at an important role for these cells in maintenance of tissue homeostasis during steady state and inflammation (Krenkel and Tacke, 2017; Mosser *et al*., 2021; Oishi and Manabe, 2018; Watanabe et al., 2019). Numerous studies have identified functionally distinct macrophage sets with polarized pro-inflammatory (M1) or anti-inflammatory (M2) phenotypes with characteristic metabolic states (Galvan-Pena and O’Neill, 2014; Tan et al., 2015; Wang et al., 2018).

With a central role of mitochondria in cellular metabolism and physiology, it is logical to presume stringent control of organelle integrity in normal as well as in response to stress. Over the last few years, infection dependent modulation of mitochondrial dynamics and function have been demonstrated as key determinants of host cell response kinetics and immune signaling (Benmoussa et al., 2018; Mehta et al., 2017; Mills *et al*., 2016; Ramond et al., 2019). A recent study has demonstrated a significant lowering of energy metabolism in macrophages on account of the strong type I IFN responses induced by Mtb infection (Olson et al., 2021). With the mitochondrial CMPK2, an ISG, one of dominantly expressed proteins in infected macrophages, we sought to investigate the importance of this gene in macrophage response dynamics in human macrophages.

We provide evidence for a critical role of the mitochondrial gene CMPK2 in regulating organelle integrity and macrophage physiology. Altered expression of this gene led to disruption of the normal mitochondrial physiology as evidenced by decreased membrane potential and increased ROS. This further translated into marked dysregulation of basal inflammation in the cells with enhanced activation of ERK signaling leading to a marked increase in production of pro-inflammatory mediators like TNFα, IL8 and IL1β. In line with a recent study of the importance of increased lactate production by M1 polarized macrophages in microbicidal capacity (Ó Maoldomhnaigh et al., 2021), both the KD and OE cells also portrayed a significant shift in their metabolic status towards rapid energy production via glycolysis and consequent accumulation of lactate in the cells. Our observation of identical phenotypic characteristics of CMPK2 silenced as well as overexpression cells (enhanced mitochondrial stress even at the basal state) hinted at the possibility of a fundamental role of this gene in organelle function and integrity. It is also interesting to assume an association of CMPK2 in a larger protein complex in cells, in which changes in expression of one component would affect the overall stoichiometric composition of the complex and its function as reported earlier in the case of SRPK1 in cancerous cells (Wang et al., 2014). The association of CMPK2 with the mitochondrial membrane in proximity to the mitochondrial ETC proteins coupled to our observed decrease in the levels of complex 1 in CMPK2 dysregulated (KD and OE) macrophages (Fig. S5), provides initial evidence for this hypothesis. The observed impairment in oxygen consumption in these macrophages and a concomitant increase of the glycolytic metabolite-lactate further supports the involvement of CMPK2 in central metabolic activities of macrophages.

We observed that the removal of MLS from CMPK2 destabilized the protein and resulted in enhanced degradation by the cellular proteolytic machinery. This observation is in sync with previous reports(Ravanelli et al., 2020) demonstrating that mitochondrial proteins without signals for mitochondrial partitioning tend to form misfolded aggregates that are rapidly degraded by the host proteasome (Wrobel et al., 2015). We also observed an absolute requirement of kinase activity for CMPK2 function and the importance of the N-terminal domain in protein stability (data not shown, part of another manuscript).

A previous study identified an important role for serine catabolism in the maintenance of cellular redox under hypoxic conditions. Repression of serine catabolism by silencing SHMT2 in cancer cells resulted in enhanced ROS under hypoxia (Ye et al., 2014). We also observed significant downregulation of genes involved in serine biosynthesis and conversion to glycine in macrophages with altered expression of CMPK2. In addition, we also see enhanced ROS and hypoxia in these macrophages. Our observation of a significant decrease in SHMT1 expression without any significant change in SHMT2 in the KD and OE macrophages highlights another possible link between serine catabolism and cellular redox maintenance via the regulated expression of CMPK2.

Serine metabolism has been shown to affect cellular physiology including inflammation in macrophages. Studies have suggested an inflammation promoting role for serine biosynthesis in macrophages with serine depletion in the culture media actively stunting the expression of pro-inflammatory genes like TNF and IL1β (Chen et al., 2020; Rodriguez et al., 2019). While these studies signify the importance of extracellular serine and its uptake by cells, intracellular serine biosynthesis and its conversion to glycine by the serine glycine and one-carbon pathway (SGOCP) genes in modulating macrophage phenotypes is yet not understood. A recent robust study has demonstrated that Phgdh mediated serine synthesis is required for expression of anti-inflammatory genes and inhibition of Phgdh resulted in enhanced expression of pro-inflammatory genes (Wilson *et al*., 2020).

To validate the PHGDH role in enhanced inflammatory gene expression, we inhibited PHGDH and observed enhanced expression of pro-inflammatory genes in macrophages. This is in concordance with a previous study by Wilson et al., wherein M2 (anti-inflammatory) macrophages showed enhanced Phgdh and Psat1 expression (Wilson *et al*., 2020). In fact, treatment with LPS that polarizes the cells to an M1 phenotype reduced Phdgh expression in macrophages while IL4 enhanced the expression again emphasizing the association of serine biosynthesis with the anti-inflammatory macrophages. Further inhibition of PHGDH resulted in heightened pro-inflammatory response as observed in the KD and OE macrophages.

Given the importance of CMPK2 in maintaining mitochondrial architecture, function and in enhancing overall inflammation, macrophage would actively induce this gene as a response to intracellular bacterial infection. With bacterial contact immediately stimulating host cell surface receptors, it is not surprising that CMPK2 would be induced in response to activation of more central TLR4 signaling that primarily leads to activation of inflammatory programs in the infected macrophages. Moreover, our data of a long-term breakdown of inflammation control in macrophages following prolonged disturbance of CMPK2 expression levels only highlights the importance of rapid and robust regulation of this gene by cells for reverting to basal status. This supports our hypothesis that CMPK2 is an important component of the cellular inflammation rheostat controlling macrophage metabolic and transcriptional response to infections. A recent study has defined the criticality of cellular pyrimidine levels in mitochondrial DNA mediated regulated innate immunity (Sprenger et al., 2021). Given the putative role of CMPK2 in the production of CTP/UTP, and the dysregulation of serine metabolism on CMPK2 imbalance, another important biosynthetic node for nucleotide biosynthesis in the cell, one can envisage a similar situation of an altered cellular *vis-a-vis* mitochondrial nucleotide levels. Comprehensive analysis towards pinpointing the precise molecular mechanism of CMPK2 mediated regulation of macrophage immuno-metabolism would help develop novel mechanisms of modulating innate response control paving the way for future host cell directed therapeutics.

## Acknowledgments

The authors thank CSIR (VR-BSC0123, MLP2012), and DBT (SG-GAP0088) funding agency for supporting the study. The authors thank Mr. Manish Kumar and CSIR-BSC0403 for the confocal microscopy facility, CSIR-STS0016 for BSL3 facility. The authors wish to thank Dr. Soumya Sinha Roy, CSIR-IGIB for the plasmid pmtDsRed. The authors thank PA’s doctoral advisory committee members for useful discussions and comments on the project. The mass spectrometry facilities are duly acknowledged. The student fellowships from CSIR: PA-CSIR-SRF, RA, PA-CSIR-BSC0124, MC-CSIR-JRF, India. The authors thank Kanika Bisht and Mahima Madan for proof reading of the manuscript.

## Author Contributions

PA, SR, SG, and VR were involved in conceptualizing and design of the work. Experiments were performed by PA, MC, TR and DS. PA and RC performed Mass spectrometry, PA, SG, and VR wrote the manuscript.

## Materials and Methods Reagents

The following chemicals were purchased from Sigma-Aldrich: U0126 (U120), RNAzol RT (R4533), Zymosan (Z4250), Thermo Fisher Scientific: MitoSOX Red (M36008), MitoTracker Deep Red (M22426) and Invivogen: LPS (tlrl-smlps), CLI095 (tlrl-cli095), poly I:C (tlrl-pic), Pam3CSK4 (tlrl-pms). Mitoquinol (89950), PHDGH-inactive (19717) and NCT-503 (19718) were purchased from Cayman Chemical Company.

### Cell culture

THP-1 cells were cultured in RPMI-1640 (Himedia laboratories, Mumbai, India) media supplemented with 10% FBS and 1mM sodium pyruvate (Himedia laboratories, Mumbai, India). HEK293T cells were cultured in DMEM supplemented with 10% FBS and 1mM sodium pyruvate. For differentiation, THP-1 cells were treated with 100nM PMA for 24hr and rested for 48hr before the next manipulation. Both the cell lines were confirmed for their identity by STR PCR analysis and maintained mycoplasma free by routine monitoring. LPS treatment of macrophages was done with 10 ng/ml LPS for 6h. TLR4 signaling was inhibited by treating with 10 μM of CLI095 before infection or LPS treatment. For ERK inhibition, THP-1 macrophages were treated with 2 μM of U0126 for 6h. To inhibit serine biosynthesis macrophages were treated with 10 μM NCT-503 for 6h and 10 μM PHGDH-inactive compound was used as negative control.

### Generation of THP1 lines with CMPK2 silencing

To generate knockdown cells, HEK293T cells were co-transfected with CMPK2 siRNA plasmid (Applied Biological Materials Inc, Richmond, Canada), packaging plasmid (pCMV-dR8.2) and envelope plasmid (pCMV-VSV-G) using Lipofectamine-LTX reagent (Thermo Fisher Scientific India Pvt. Ltd, Mumbai, India). After 6h of transfection, media was removed, and fresh media was added to the cells. After 48h of transfection, the culture supernatant was collected, centrifuged at 500 rpm to remove cell debris. The virus particles in the supernatant were concentrated to 100μl using Amicon® Ultra-15 Centrifugal Filters (Sigma-Aldrich Chemicals Private Limited, Bengaluru, India) and used to infect THP1 cells. Cells were then selected with 0.6 μg/ml puromycin and GFP positive clones were used for further studies.

### Bacterial culturing and infection

*Mycobacterium tuberculosis* strain Erdman was grown in 7H9 Middlebrook media BD Biosciences, USA) supplemented with Middlebrook ADC (BD, USA) at 37^°^C. *E. coli and Salmonella enterica serovar Typhimurium* were grown in Luria-Bertani (LB) media (Himedia Laboratories, Mumbai, India) at 37°C. Single cell suspension of mid-log phase bacteria was adjusted to the required cell density and then used to infect the differentiated THP-1 macrophages at multiplicity of infection (MOI) of 5. After 6h *p*.*i*. for Mtb, cells were washed with PBS to remove extracellular bacteria, and at various times post infection, bacterial numbers were enumerated by lysis of cells and plating for CFU in 7H10 agar plates. For *S. typhimurium*/ *E. coli*, post addition of bacteria at a MOI of 10, the plates were centrifuged for 5 min and then left for 20 min at 37^º^C. The media was removed and treated with 100μg/ml of gentamicin for 2h in RPMI at 37^º^C. The cells were washed with PBS for 3 times, and infection was continued in media containing 12μg/ml of gentamicin and at indicated time points cells were lysed with PBS-Triton-X-100 (1%) for CFU plating on LB agar (Himedia laboratories, Mumbai, India) plates. The colonies were counted and is represented as CFU/well.

### Gene expression analysis by qPCR

Total RNA was isolated using RNAzol (Sigma-Aldrich, USA) method and the concentration was quantified using Nanodrop 2000 UV-visible spectrophotometer. cDNA was prepared with 250-1000 ng total RNA by a RT-PCR using Verso cDNA synthesis kit (Thermo Fisher Scientific, USA). qPCR was performed on a Roche LC480 II system using DyNAmo Flash SYBR Green mix (Thermo Fisher Scientific, USA). Primer sequences are given in Table 1.

**Table 1:**
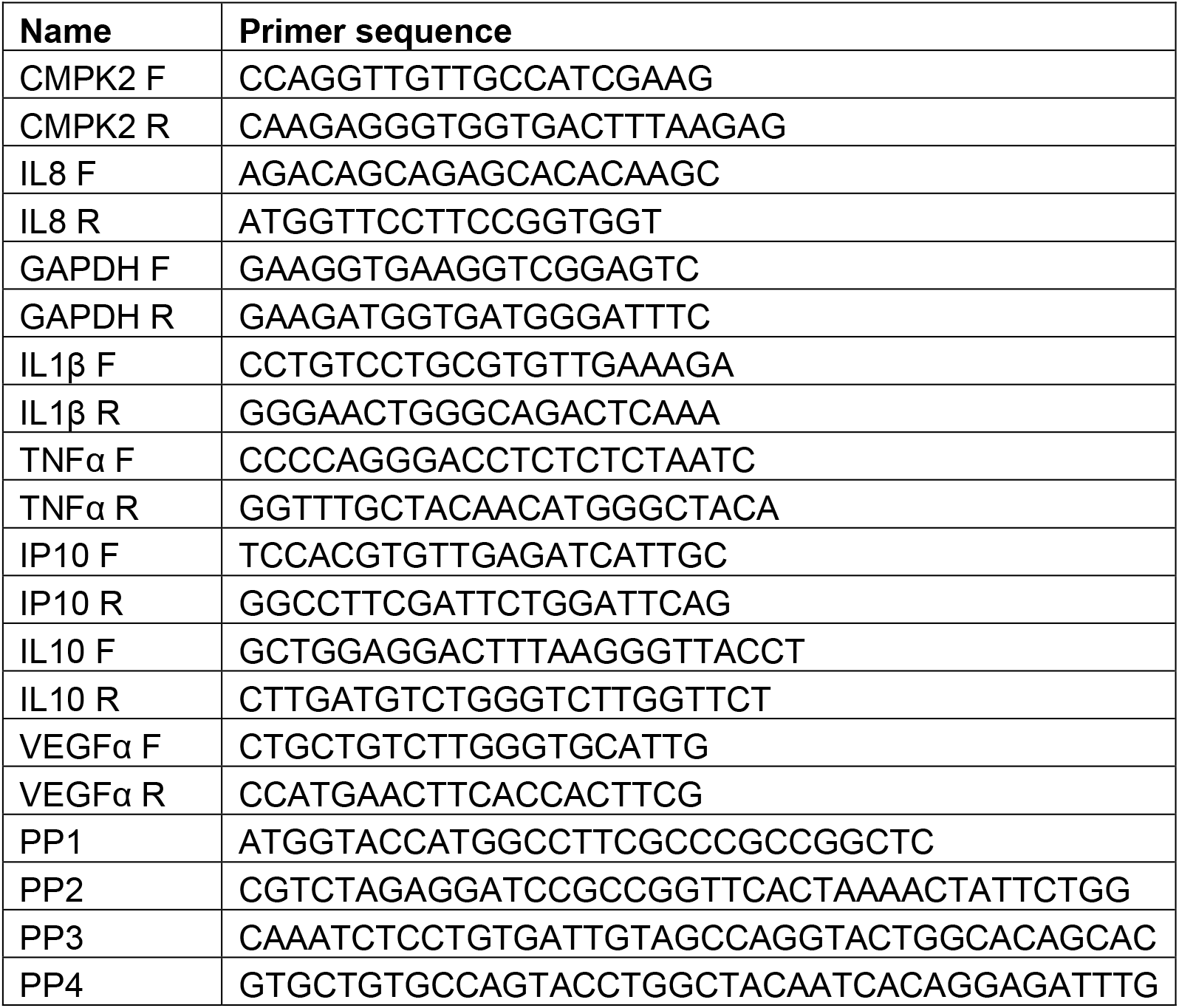
List of primers used in this study.

### Cloning of CMPK2 and the D330 variant

The NFkB promoter from pNFkB-d2GFP was excised by digestion with *Ecl*136II and *Hind*III and the mCherry fragment from pmCherry-N1was removed with *Hind*III and *Hpa*I and ligated to *Nru*I and *Eco*RV digested pcDNA3.1(+) to get the plasmid pPRAM1. The NFkB promoter from pPRAM1 was replaced with EF-1α from pTracer-EF/V5 His A by using *Mlu*I and *Eco*RI to get pPRAM2. The CMPK2 fragments were amplified from THP1 cDNA using the primer sets PP1 and PP2 and cloned into the pPRAM2 using *Kpn*I and *Bam*HI. The D330A catalytic site mutant was prepared by site directed mutagenesis using the primers PP3 and PP4. The sequences were verified by sequencing. The list of primers used in this study is given in table 1.

### Western blotting and ELISA

For immunoblotting, cells were lysed in RIPA buffer, resolved by SDS-PAGE and transferred to nitrocellulose membrane. After blocking with 5% BSA in Tris Buffered Saline Tween (TBST) for 1hr at room temperature, membranes were incubated with the appropriate dilution of the primary antibody overnight, washed with PBS and probed with infrared dye conjugated secondary antibody and developed in the LI-COR Odyssey platform. The following antibodies were purchased from Cell Signaling Technology: Rabbit anti-α-tubulin (2125), Rabbit anti-phospho-p42/44 (4377), Rabbit anti-p42/44 (4695), Rabbit anti-phospho-NFkB p65 (3033), Rabbit anti-NFkB p65 (4764), Rabbit anti-phospho-JNK (4671), Rabbit anti-JNK (9258), Abcam: Rabbit anti-mCherry (ab167453), Mouse anti-TOM20 (ab56783), Rabbit anti-TIM50 (ab109436), Mouse anti-MFN1/2 (ab57602), Mouse anti-OPA1 (ab194830), Rabbit anti-DRP1 (ab140494), Rabbit anti-HIF1α (ab51608) BD Bioscience: Rabbit anti-VDAC (D73D12), Mouse anti-SOD2 (611581) and LI-COR Biotechnology: Goat anti-Rabbit IgG 800 (926-32211), Goat anti-Mouse IgG 680 (926-68070). Rabbit anti-CMPK2 (HPA041430) was purchased from Sigma-Aldrich.

For ELISA, either cell supernatants or cell extracts were used for cytokine estimation with specific kits-TNFα (cat no: 88-7346-77, eBioscience) and IL1β (cat no: 557953, BD Biosciences) ELISA kits according to manufacturer’s protocol.

### Analysis of ROS by FACS

THP1 monocyte derived macrophages (MDM) were removed with PBS and 4mM EDTA, washed with PBS, resuspended in RPMI media and used for staining with 5μM MitoSOX Red and 100nM MitoTracker Deep Red at 37°C for 30 mins. Cells were then washed twice in HBSS and finally resuspended in 500μl HBSS and analyzed in FACS ARIA II (BD Biosciences).

### Metabolite measurement

Intracellular metabolites for MS-based targeted metabolomics were extracted using methanol-water. Briefly, 1.2 × 10^6^ cells from each condition were washed three times with ice-cold PBS, and then quenched with methanol-water (4:1). The cell suspension was freeze thawed in liquid nitrogen three times. The suspension was centrifuged at 15,000 g at 4°C for 10 min. Supernatant was collected, vacuum dried and then reconstituted in 50 μl of 50% methanol. The reconstituted mixture was centrifuged at 15,000 g for 10 min, and 5 μl was injected for LC–MS/MS analysis.

The data were acquired using a Sciex Exion LCTM analytical UHPLC system coupled with a triple quadrupole hybrid ion trap mass spectrometer (QTrap 6500; Sciex) in negative ion mode. Samples were loaded onto an Acquity UPLC BEH HILIC (1.7 μm, 2.1 × 100 mm) column, with a flow rate of 0.3 ml/min. The mobile phases comprising of 10 mM ammonium acetate and 0.1% formic acid (buffer A) and 95% acetonitrile with 5 mM ammonium acetate and 0.1% formic acid (buffer B). The linear mobile phase was applied from 95% to 20% of buffer A. The gradient program was used as follows: 95% buffer B for 1.5 min, 80%–50% buffer B in next 0.5 min, followed by 50% buffer B for next 2 min, and then decreased to 20% buffer B in next 50 s, 20% buffer B for next 2 min, and finally again 95% buffer B for next 2 min. After data acquisition, peaks corresponding to each metabolite were extracted using Sciex Multiquant TM v.3.0 software and the area was exported in an excel sheet. Normalization was performed using total area sum of that particular run.

### Oxygen Consumption

3-4 × 10^6^ cells of THP1 MDMs in complete RPMI were added to the chambers of the Oxygraphy-2k (O2k, Oroboros Instruments, Innsbruck, Austria). The oxygen concentration was measured over time at 37ºC under constant stirring. The oxygen consumption rate was calculated using XY graph of oxygen consumption and time.

### Transcriptome analysis

Total RNA was isolated using RNAzol (Sigma Aldrich, India) and transcript sequencing was done commercially by Bencos research solutions private ltd., (Bengaluru, India). cDNA libraries were generated using Truseq RNA Library Prep Kit (Illumina San Diego, USA) and sequenced in an Illumina Novaseq 6000 platform. RNA-seq reads were aligned with hg38 genome using STAR (Dobin et al., 2013). HTseq-count was used to count the transcripts (Anders et al., 2015). The differential expression analysis across samples was analyzed using GSEA (Subramanian et al., 2005).

### Confocal Microscopy and Image analysis of stained cells

THP1 monocytes (stable transfectants of non-targeting or CMPK2 specific siRNA) were transfected with plasmid DNA mtDsRed (kind gift by Dr. Soumya Sinha Roy, CSIR-IGIB) and selected for stably expressing clones with 400μg/ml of G418. Cells were differentiated on coverslips, fixed with 4% formaldehyde and imaged in a Leica 480 confocal microscopy. For live imaging, monocytes overexpressing CMPK2 were plated at a cell density of 0.3 × 10^6^ cells/ml in the cell culture dish containing two chambers per dish (1 ml/ chamber). Mitotracker Green dye was added to the media and the cells were kept in 37ºC CO_2_ incubator for 20mins. The cells were imaged on a Leica SP8 confocal microscope.

### Mitochondrial localization

Differential centrifugation was employed for mitochondria isolation from the cells (Frezza et al., 2007). Briefly, 2.5 × 10^6^ THP1 MDMs were washed twice with PBS and resuspended in 3 ml ice-cold cell isolation buffer (IBc-containing 10mM Tris pH to 7.4, 1mM EGTA pH to 7.4 and 0.2M sucrose). The cells were homogenized using a glass Teflon pestle potter. The homogenate was then centrifuged at 600g for 10 min at 4ºC, the supernatant was collected and centrifuged at 7000g for 10 min at 4ºC. The pellet was then washed and resuspended in 200 μl IB_C_ and transferred to a 1.5ml Eppendorf tubes. The homogenate was again centrifuged at 7000g for 10 min at 4ºC. The supernatant was discarded and the pellet was resuspended to get the mitochondrial fraction.

For Proteinase K digestion, the mitochondria were resuspended in MS buffer (210mM mannitol, 70mM sucrose, 5mM Tris-HCL-pH 7.5, 1mM EDTA-pH 7.6)/ 20mMHEPES with or without 2 mg/ml Proteinase K (for 15 min) and/or 1% Triton-X-100. The reaction was terminated by the addition of 5 mM phenylmethylsulfonyl fluoride. The mixture was centrifuged at 15000g for 15 min and the pellet fraction was collected. For analysis of membrane proteins, the mitochondrial fraction was resuspended in MS buffer with and without 0.1M Na_2_CO_3_ and incubated in ice for 30 min. The pellet and supernatant fractions were collected at 15000g for 15 min. All fractions were analyzed by immunoblotting with respective antibodies.

### Graphs and Statistical analysis

Statistical analyses were performed by using the two tailed Student’s t-test. GraphPad Prism software was used for graphs and statistical analysis.

### DATA availability statement

The RAW data of RNA-seq experiments of this study have been uploaded in NCBI-GEO with accession number -GSE199951.

## Supplementary Figures

**Fig. S1:**
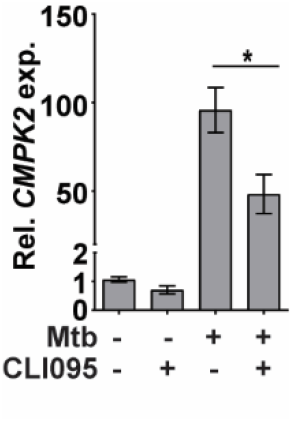
Expression of *CMPK2* in THP1 macrophages upon Mtb infection in the presence or absence of CLI095.

**Fig. S2:**
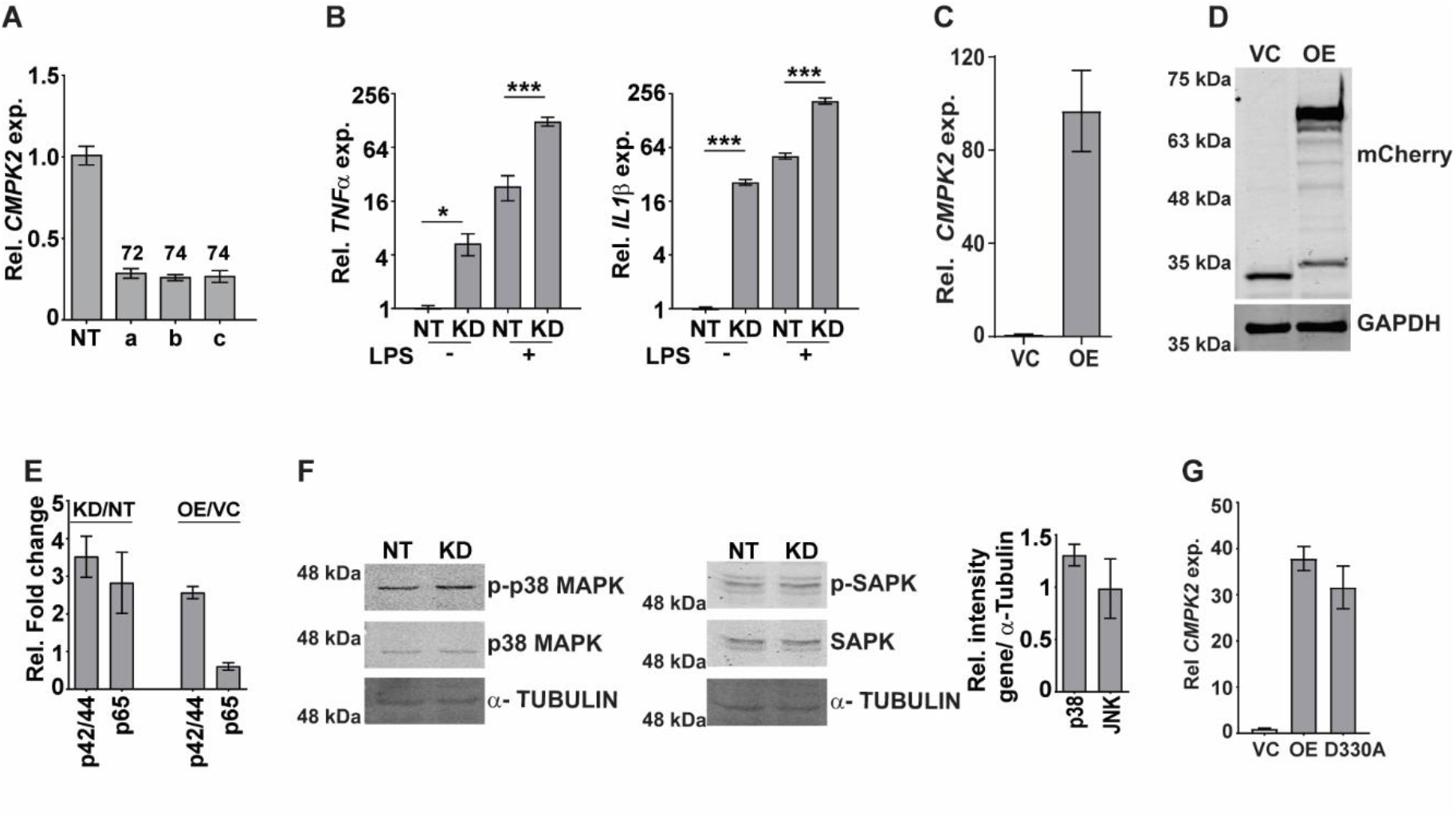
[A] Expression of *CMPK2* in THP1 macrophages stably expressing the non-targeting (NT) siRNA or three different siRNAs (a-c) against CMPK2 by qPCR. The gene expression was normalized with *GAPDH* and relative fold changes compared to NT are represented as mean + SEM (N=2). [B] Expression of *TNFα* and *IL1β* gene in NT and KD macrophages following LPS stimulation for 6h. The relative gene expression folds in triplicate assay wells are represented with respect to *GAPDH* as mean + SEM for N=2. [C] Expression of *CMPK2* in OE or empty vector -VC cells was analyzed by qPCR. The relative gene expression folds in triplicate assay wells are represented with respect to *GAPDH* as mean + SEM for N=3. [D] CMPK2 expression in VC and OE cells was analyzed by immunoblotting with tag specific (mCherry) antibody. [E] Relative quantitation of p-ERK and p-NFκB in the NT, KD, VC, OE cells by densitometric analysis of immunoreactivity is shown. Values are mean + SEM of triplicate (N=3) independent blots. [F] Analysis of activation of the p38 MAPK and SAPK/JNK signaling pathways in NT and KD macrophages by immunoblotting with antibodies specific for the phosphorylated (active) and non-phosphorylated forms of the proteins. The relative intensities of the blots in the CMPK2 silenced cells w.r.t control (NT) cells ar e represented as mean + SEM of N=3. Expression of α-TUBULIN was used as control. [G] Expression of *CMPK2* in VC, OE and D330A cells was analyzed by qPCR and is depicted relative to GAPDH levels of N=3 assays.

**Fig. S3:**
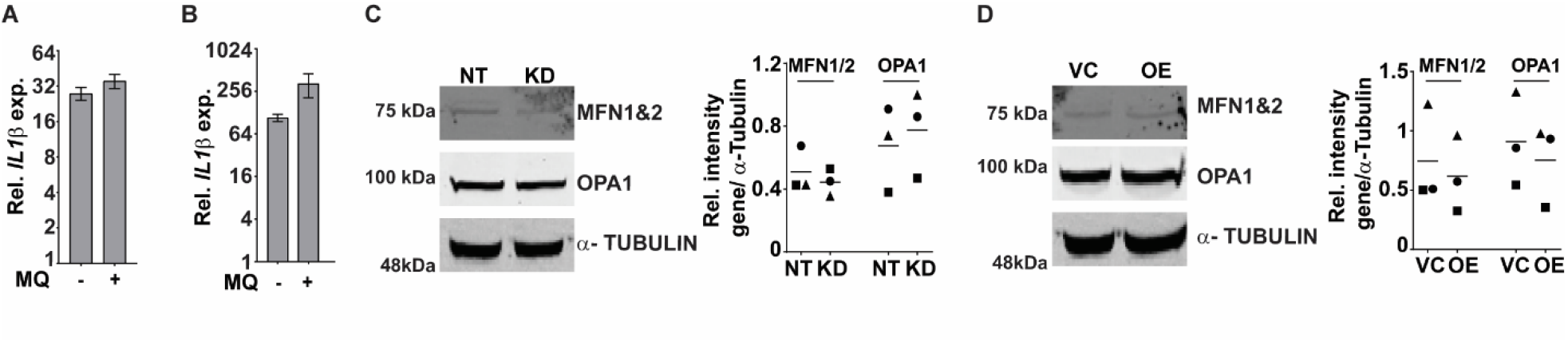
[A, B] Expression of *IL1β* in the macrophages with the addition of a specific mitochondrial ROS inhibitor-MQ at day 2 [A] and day 3 [B] of PMA treatment intervals post activation was analyzed by qPCR. Values are mean fold change in expression with respect to *GAPDH* + SEM for triplicate assays of N=2/3 experiments. [C, D] Expression of proteins involved in mitochondrial fusion by immunoblotting with specific antibodies. MFN1&2 and OPA1 were checked in NT, KD [C] or VC, OE [D] cells along with α-TUBULIN levels as a control. Relative intensity values are depicted + SEM of N=3.

**Fig. S4:**
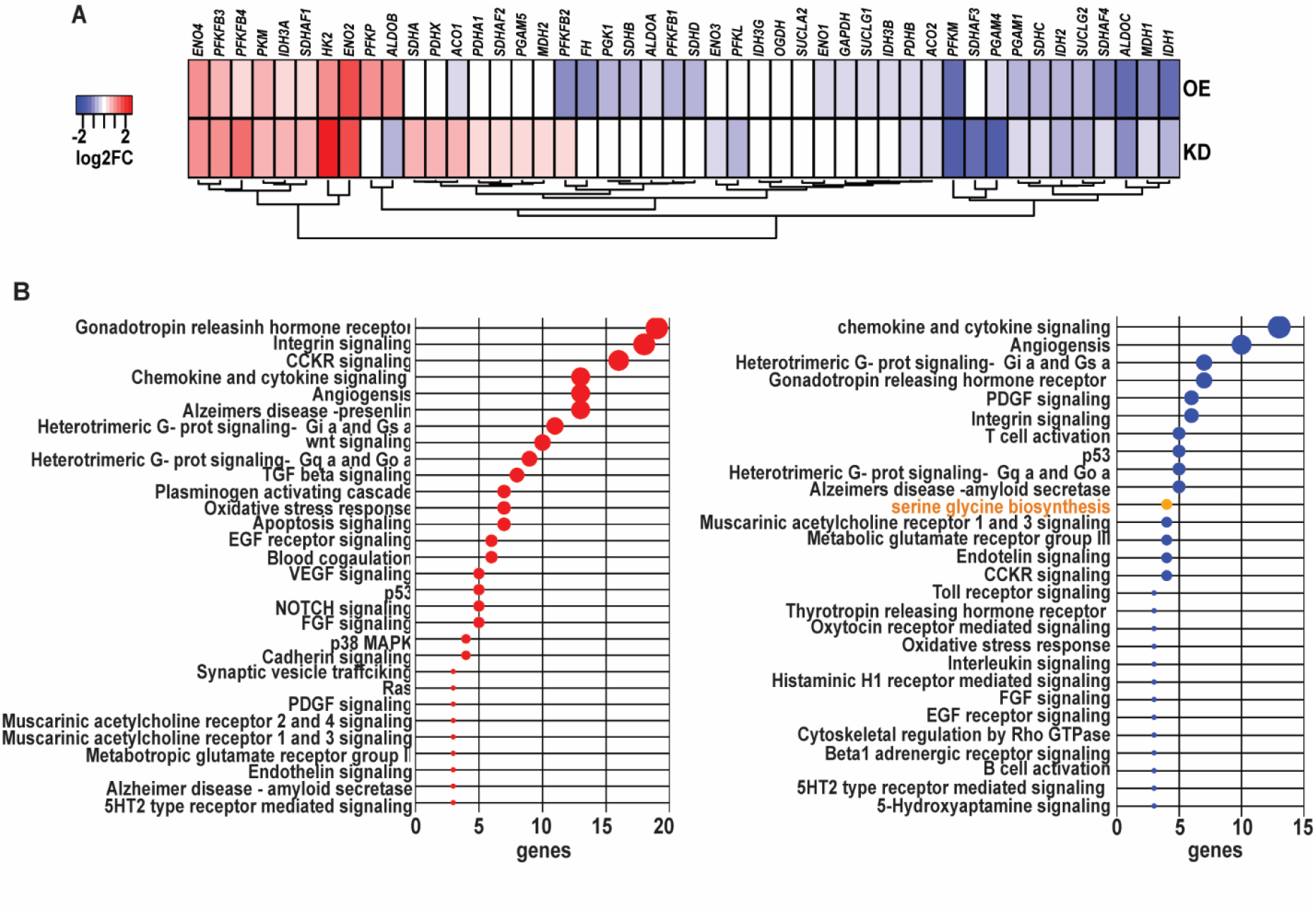
[A] Expression of genes involved in glycolysis and TCA was represented as heatmap. The expression was extracted from the RNA sequencing data of CMPK2 dysregulated THP1 cells. The values are represented as log_2_ compared to respective control cells. [B] Panther gene family enrichment analysis of the commonly up and down regulated genes in the KD and OE macrophages is represented as a bubble plot. X axis is the number of genes of the pathway.

**Fig. S5:**
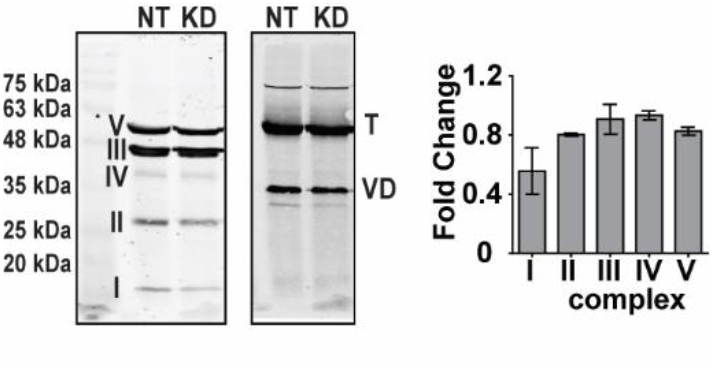
Analysis of expression of protein complexes of the mitochondrial respiratory chain in cell lysates of NT and KD macrophages by immunoblotting. α-TUBULIN (T) and VDAC1 (VD) were used as loading controls. The band intensities of the complex proteins were normalized to α-TUBULIN and the relative intensity of KD w.r.t. NT is represented as fold change + SEM for N=2.

